# Monocyte-derived Prostaglandin E2 inhibits antigen-specific cutaneous immunity during ageing

**DOI:** 10.1101/2020.04.02.020081

**Authors:** Emma S Chambers, Milica Vukmanovic-Stejic, Barbara B Shih, Hugh Trahair, Priya Subramanian, Oliver P Devine, James Glanville, Derek Gilroy, Malcom Rustin, Tom C Freeman, Neil A Mabbot, Arne N Akbar

## Abstract

Ageing results in a decline in immune function. We showed previously that healthy older humans (>65 years old) have reduced antigen-specific cutaneous immunity to varicella zoster virus (VZV) antigen challenge. This was associated with p38 MAP kinase driven inflammation that was induced by mild tissue injury caused by the injection of the antigen itself. Here we show that non-specific injury induced by injection of air or saline into the skin of older adults recruits CCR2^+^CD14^+^ monocytes by CCL2 produced by senescent fibroblasts. These monocytes reduced T_RM_ proliferation via secretion of prostaglandin E2 (PGE_2_). Pre-treatment with a p38-MAPK inhibitor (Losmapimod) in older adults *in vivo* significantly decreased CCL2 expression, recruitment of monocyte into the skin, COX2 expression and PGE_2_ production. This enhanced the VZV response in the skin. Therefore, local inflammation arising from interaction between senescent cells and monocytes leads to immune decline in the skin during ageing, a process that can be reversed.

**Summary:** Inflammation resulting from tissue injury blocks antigen-specific cutaneous immunity during ageing. Monocytes recruited to the skin inhibit T_RM_ function through COX2-derived prostaglandin E_2_ production. Blocking inflammation and resulting prostaglandin E_2_ production with a p38-MAP kinase inhibitor significantly enhances cutaneous antigen-specific responses.

## Introduction

Immunity decreases during ageing as evidenced by the increase in susceptibility to infections such as pneumonia and influenza, re-activation of latent infections such as varicella zoster virus (VZV; herpes zoster or shingles), decreased vaccine efficacy and increased incidence of cancer (Ciabattini et al., 2018; Diffey and Langtry, 2005; Gavazzi and Krause, 2002). A key challenge is to identify mechanisms that can be manipulated to boost immunity and reduce morbidity and mortality in older individuals. Low grade systemic inflammation, termed inflammageing is characterised by high serum levels of the inflammatory cytokines IL-6, IL-1β, TNFα and C reactive protein (CRP) (Franceschi et al., 2017). Elevated levels of inflammatory proteins are a strong predictor for frailty and mortality (Dinh et al., 2019; Furman et al., 2017). However, it is not clear how inflammageing inhibits antigen-specific immunity during ageing. Mechanisms that contribute to inflammageing may include increased gut permeability and subsequent lipopolysaccharide (LPS; a TLR4 ligand) leakage (Kim et al., 2016; Thevaranjan et al., 2017), increased presence of damage-associated molecular pattern molecules (DAMPs) and increased visceral fat that can activate mononuclear phagocytes (Franceschi et al., 2017). In addition, senescent non-lymphoid cells in tissues (such as fibroblasts) can also secrete many inflammatory proteins as part of the senescence associated secretory phenotype (SASP) (Campisi, 2013; Freund et al., 2010; Hall et al., 2016).

We previously identified reduced antigen-specific cutaneous recall responses in healthy old compared to young donors *in vivo* (Agius et al., 2009; Akbar et al., 2013; Vukmanovic-Stejic et al., 2008; Vukmanovic-Stejic et al., 2018). This difference was not due to decreased numbers or the responsiveness of antigen-specific circulating or skin resident-memory T cells (T_RM_) (Vukmanovic-Stejic et al., 2015), suggesting that there were changes in the skin microenvironment that inhibited antigen-specific T cell function in these subjects. One such change was the propensity to exhibit amplified inflammation in response to mild tissue injury in the skin after saline-injection, that was not observed in young subjects (Vukmanovic-Stejic et al., 2018). There was a direct association between the propensity to exhibit inflammation in older subjects and the decreased response to VZV antigen challenge in the skin since short-term systemic treatment with an oral anti-inflammatory drug (Losmapimod; p38 mitogen-activated protein kinase [p38 MAPK] inhibitor) significantly increased the cutaneous VZV response (Vukmanovic-Stejic et al., 2018). A crucial unanswered question was the mechanism by which inflammation inhibited immunity *in vivo*.

Here we show that CCR2^+^CD14^+^ monocytes are recruited into the skin of older donors after saline, air or VZV antigen injection. These monocytes are recruited by in response to CCL2 secreted by senescent dermal fibroblasts. The infiltrating monocytes have increased expression of cyclooxygenase 2 (COX2) and can inhibit T_RM_ proliferation *in vitro* via production of prostaglandin E2 (PGE_2_). Blockade of systemic inflammation using a p38 MAPK inhibitor (Losmapimod) in older adults significantly reduced CCL2 production from senescent fibroblasts and recruitment of monocytes into the skin after injection *in vivo*. In addition, Losmapimod treatment significantly reduced COX2 expression and PGE_2_ production confirming that p38 MAPK signalling is upstream of COX2 (Dean et al., 1999; Guan et al., 1998). This significantly enhanced VZV-specific T cell proliferation in the skin. This identifies a broad mechanism by which inflammation may inhibit immunity in many different tissues during ageing. A key observation is that this decline in immune responsiveness may be reversible in humans by the shot-term blockade of systemic inflammation *in vivo*.

## Materials and Methods

### Study design

The investigation of healthy young versus old subjects was approved by the Ethics Committee Queen’s Square (London) and by institutional review board (UCL R&D). Healthy young (<40 years) and old (>65 years; see supplementary Table 1 for donor characteristics) were recruited for the study and individuals with history of neoplasia, immunosuppressive disorders or inflammatory skin disorders were excluded as described in detail elsewhere (Vukmanovic-Stejic et al., 2018). In brief, we excluded individuals with co-morbidities that are associated with significant internal organ or immune dysfunction including heart failure, severe COPD, diabetes mellitus and rheumatoid arthritis and individuals on immunosuppressive regimes for the treatment of autoimmune or chronic inflammatory diseases (e.g. oral glucocorticoids, methotrexate, azathioprine and cyclosporin). In addition, individuals with history of liver disease or elevated liver transaminases (>1.5 times the upper limit of normal), or who had abnormal ECG. Only Caucasian European individuals were included in the study. All volunteers provided written informed consent and study procedures were performed in accordance with the principles of the declaration of Helsinki.

For the study involving Losmapimod, we recruited 42 healthy older adults (>65 years); 19 males and 23 females with a mean age of 70.7 years old (95% CI 69-72.3 years old). VZV skin test was performed and biopsies collected at different time points as described (Vukmanovic-Stejic et al., 2018). Any individual with a clinical score >3 was excluded from the study as described previously (Vukmanovic-Stejic et al., 2018).

Two to three months after the first VZV skin challenge the same volunteers received 15 mg Losmapimod (GW856553) BID for 4 days (provided by Glaxo-Smith-Klein under a Medical Research Council Industrial Collaboration Agreement). In preliminary experiments we found that the first VZV challenge did not significantly boost the response to re-challenge of the same individuals after 2-3 months (Vukmanovic-Stejic et al., 2018). Losmapimod 15 mg BID dose used in this study was chosen on the basis of the PK, PD and safety profiles of Losmapimod observed in GSK Phase I and II studies (Watz et al., 2014). These individuals were then re-challenged with VZV skin antigen test and biopsies were taken at the same timepoints as before Losmapimod treatment. Clinical scores after VZV injection were performed by two independent researchers and an average reading was recorded. Serum CRP levels were measured with a high sensitivity assay (pre- and post-Losmapimod). No adverse side effects were observed in our study group in response to four-day Losmapimod treatment.

### Skin tests and punch biopsy

VZV antigen (BIKEN, The Research Foundation for Microbial Diseases of Osaka University, Japan) or 0.9% saline solution or an equal volume of air was injected intradermally into sun unexposed skin of the medial proximal volar forearm as per manufacturer’s instructions. Induration, palpability and the change in erythema from baseline were measured and scored on day 2 or 3 as described previously (Akbar et al., 2013). A clinical score (range 0-10) based on the summation of these parameters was then calculated. Donor characteristics can be found in Table 1.

Punch biopsies (5 mm diameter) were collected from the injection site from normal (unmanipulated skin) at various time-points (as indicated). For Losmapimod studies biopsies were collected for two time points from separate arms. Biopsies were frozen in OCT (optimal cutting temperature compound; Bright Instrument Company Ltd, Luton, U.K.). 6μm sections were cut and left to dry overnight and then fixed in ethanol and acetone and stored at -80°C.

### Immunofluorescence

Sections were stained with optimal dilutions of primary antibodies and followed by relevant secondary antibody conjugated to various fluorochromes as described. Antibodies used include CCL2 (ab7814/ab9669), CD62E (ab36688), COX2 (ab23672), EP4 (ab92763), FSP-1 (S100A4; ab93283/ab93283) from Abcam (Cambridge, U.K.), CD4 (YNB46.1.8), CD8 (YTC141.1HL) from AbD Serotec (Oxford, U.K.), CD69 (FN50) from Biolegend (London, U.K.), CD31 FITC and Ki67 FITC from BD Biosciences (Oxford, U.K.), p16 (CDKN2A; 5A8A4) Sigma Aldrich (Gillingham, U.K.) and CCR2 (PA5-23037) from Thermo Fisher Scientific (Loughborough, U.K.). Skin sections were imaged on the AxioScan Z1 slide scanner and analysed using Zen Blue (Zeiss, Cambridge U.K.) For normal and 6 hour injected skin and endothelium counts, total cells in the upper dermis were counted and numbers of cells per mm^2^ were calculated. Images were taken from the upper dermis, an example of which is shown in Figure 1. For VZV injected skin (days 2-7) the number of cells in five of the largest perivascular infiltrates present in the upper and mid dermis were selected for analysis and an average was calculated as described previously (Vukmanovic-Stejic et al., 2008).

**Figure 1:**
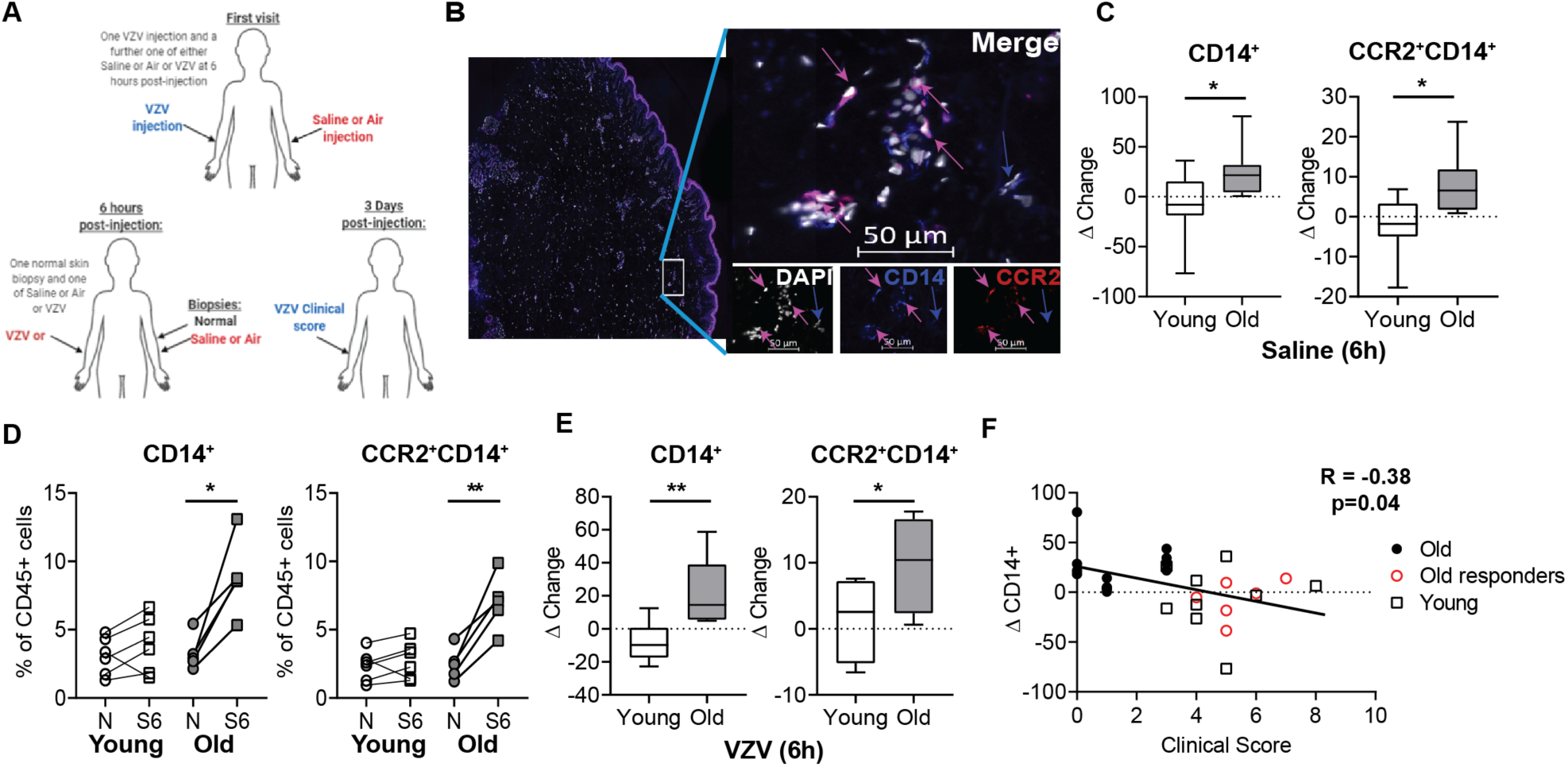
Recruitment of monocytes to sites of injection in the skin. **A**, Schematic of visits for blood and skin sampling by old (>65 years) and young (<40 years) donors taking part in this study. **B**, representative image of a skin section stained with DAPI (white), CCR2 (red) and CD14 (blue), pink arrows indicate CD14 and CCR2 co-staining and blue arrow indicates CD14 single stain. Cumulative delta change in number of CD14+ or CCR2^+^CD14^+^ cells in **C**, saline-injected (10 young and 14 old). **D**, cumulative data showing the frequency of monocytes in normal (N) and Saline (6 hours; S6) in skin biopsies collected from young (white) and old (grey) individuals as assessed by flow cytometry. Cells were identified as mononuclear phagocytes if they were CD45^+^Lineage positive but negative for CD3, CD19, CD20 CD56 and HLA-DR^+^. The CD14^+^ and CD14^+^CCR2^+^ cells were assessed as a percentage of CD45+ cells present in the skin. **E**, Cumulative delta change in number of CD14^+^ or CCR2^+^CD14^+^ cells in VZV-injected skin a (5 Young and 5 old) as compared to normal skin. **F**, Correlation between delta change in number of CD14^+^ cells in the skin after saline injection as compared to normal skin in relation to their VZV clinical response. Old donors are in black filled circles (VZV score 0-3), Old responders in open red circles (VZV clinical score ≥4) and young donors are in open black square. **C-E** analysed with an unpaired t-test and **F** analysed by Spearmans correlation. *= p<0.05; **= p<0.01.

### Skin Biopsy digestion for flow cytometric analysis

Skin biopsies (5 mm) were taken from normal and saline injected skin (6 or 24-hours post-injection) and disaggregated by overnight incubation (37°C) in either 0.8 mg/ml collagenase IV (Sigma Aldrich) with 20% FCS or a whole skin digestion kit from Miltenyi Biotec (Cambridge, U.K.). Single cell suspensions were obtained and subsequently stained with the following antibodies against: CD14 (HCD14), CD16 (3G8), CD19 (HIB19), CD20 (2H7), CD56 (HCD56), CD163 (GHI/61) and Zombie Green Fixable Viability Kit (Biolegend) and HLA-DR (G46-6), CD3 (UCHT1) and CD45 (2D1) (BD Biosciences). Samples were assessed by flow cytometric analysis on a BD Fortessa using FACSDIVA software, and subsequently analysed using FlowJo Version X (BD Biosciences).

### RNAseq analysis of skin

Three 3 mm punch biopsies were collected from each volunteer: the VZV injection site at 48 hours post-injection, saline injection site at 6 hours post-injection and normal (un-injected) skin as a control. Biopsies were immediately frozen in RNAlater. Frozen tissue was homogenized, and total RNA was extracted from bulk tissue homogenates using RNeasy Mini Kit (Qiagen). Library preparation for RNAseq was performed using the Kappa Hyperprep kit (Roche Diagnostics) and sequencing was performed by the Pathogens Genomic Unit (UCL) on the Illumina Nextseq 500 (Illumina) using the NextSeq 500/550 High Output 75 cycle kit (Illumina) according to manufacturers’ instructions, giving 15-20 million 41bp paired end reads per sample. Reads were aligned to Genome Reference Consortium Human Build 38 (GRCh38) using Hisat2 (Kim et al., 2015). Samtools was used to select for reads with paired mates. Transcript assembly was carried out using StringTie (Pertea et al., 2015), with gene-level Fragments per Kilobase of transcript per Million mapped read (FPKM) generated using Ballgown (Frazee et al., 2015). The RNAseq data has been deposited on Gene Expression Omnibus (Accession Number: GSE130633).

Statistical comparisons were made on gene count estimates generated by StringTie. Genes with low expression (<1 count per million mapped reads in 95% of the samples) or short transcript lengths (<200 nucleotides for the longest transcript) were removed. The count matrix was normalised using the TMM method in edgeR (version 3.22.5) (Robinson et al., 2010), followed by contrast fit with voom in limma (version 3.36.5) (Ritchie et al., 2015), treating the subject ID as a blocking variable. Genes with an adjusted p-value of less than 0.05 and expression change of greater than 2-fold up or down, were considered to be statistically significant.

Gene correlation networks (GCN) were constructed and visualised using Graphia Professional (Kajeka Ltd, version 2.1). Genes without a known HUGO gene nomenclature, or with expression < 1 FPKM for all samples, were removed from the co-expression network analysis. Markov Cluster Algorithm (MCL) was used for defining network clusters using an inflation value of 2.2.

### Leukocyte isolation from peripheral blood

PBMC were isolated by density centrifugation using Ficoll-Paque (Amersham Biosciences, Little Chalfont, Buckinghamshire, UK). CD4^+^ T cells were purified by positive selection and monocytes were isolated by negative selection according to the manufacturer’s instructions (Miltenyi Biotec, Woking, UK).

### Monocyte and T cells co-culture

Monocytes (1×10^6^/ml) were cultured in the presence of LPS (1ng/ml) and/or NS-398 (1µM) at 37°C with 5% CO_2_. Monocytes were collected three hours post incubation and washed once to remove residual LPS. The monocytes were pre-incubated with inhibitors NS-398 (1µM) or Losmapimod (3µM) where indicated. CD4^+^ T cells were labelled with CellTrace Violet (Invitrogen) according to the manufacturer’s instructions prior to culture. CD4^+^ and monocytes (unstimulated or LPS stimulated) were co-cultured at different ratios, in the presence of plate-bound CD3 (1µ/ml) and IL-2 (50IU/ml) and MF-498 (1µM) as indicated (at 37°C with 5% CO_2_) for four days.

For experiments involving co-culture of monocytes with skin T_RM_ cells, two suction blisters were formed over uninjected skin, as described previously (Akbar et al., 2013). Cells were collected from the blister the following day and incubated on plastic for 3 hours, to remove mononuclear phagocytes. Cell suspensions (predominantly CD4^+^ T cells) were removed and labelled with CellTrace Violet and co-cultured with unstimulated or LPS stimulated monocytes as described above.

### T cell proliferation assessment

Samples were collected at day 4 post-culture and stained for the following: CD3 (UCHT1), CD4 (RPA-T4; BD Biosciences) and Zombie NIR™ Fixable Viability Kit (Biolegend). Samples were assessed by flow cytometric analysis on a BD Fortessa using FACSDIVA software, and subsequently analysed using FlowJo Version X (BD Biosciences).

### Western Blot analysis

Cell pellets from 3 hour monocyte co-cultures were harvested and lysed with RIPA buffer (Sigma-Aldrich) supplemented with protease and phosphatase inhibitors (Cell Signalling) for 30 min on ice. Cell lysates were diluted in SDS sample buffer with reducing agent (NuPage, Life Technologies) and boiled for 5 min at 95 °C. Samples were separated by protein electrophoresis at 120 V for 2 h using 10% Bis-Tris pre-cast gels (NuPage) and transferred overnight at 4°C onto Hybond-P PVDF membranes (GE Healthcare). After blocking in ECL blocking agent (GE Healthcare), membranes were probed with primary antibodies overnight at 4°C. Primary antibodies used were COX2 (ab15191; Abcam), phospho-p38 MAPK (T180/Y182; 9211), and p38 MAP Kinase (9212), GAPDH (2118; Cell Signalling). The membrane was washed and incubated with HRP-conjugated secondary antibodies (GE Healthcare, 1:4000) for 1 h at room temperature. Antibodies were detected using the ECL detection kit (GE Healthcare). Prior to re-probing with different antibodies, membranes were stripped at room temperature with agitation using Restore stripping buffer (Thermo Scientific). Protein bands were quantified using ImageJ software. The integrated density of each band was normalised to GAPDH using the gel analysis function of ImageJ.

### Dermal Fibroblast isolation and culture

Dermal fibroblasts were isolated and cultured as described previously (Pereira et al., 2019). senescence was induced in primary human dermal fibroblasts by exposure to X-ray radiation at a total dose of 10Gy at a rate of 5Gy/min. Irradiated cells were cultured for a further 10-28 days to allow for senescence to develop. Senescence was confirmed by staining for SAβ-galactosidase (Cell signalling, London U.K.). Non-senescent (early passage number 4-10) and senescent fibroblasts were cultured in a 48-well plate at 20 × 10^4^ cells per well in the presence of absence of Losmapimod (3µM), and supernatants were collected at 3 and 24 hours.

### Assessment of supernatants

Cytokine concentration in culture supernatants were assessed by cytometric bead array (CBA) according to the manufacturer’s protocol. Samples were analysed using a BD Verse flow cytometer (BD Biosciences). The lower limit of detection for each analyte was 1.5pg/mL. For assessment of PGE_2_ production from monocyte cultures with a Prostaglandin E_2_Direct Biotrak Assay used according to the manufacturers protocol (Fisher scientific).

### Statistics

Statistical analysis was performed using GraphPad Prism version 8.00 (GraphPad Software, San Diego, California, USA). Data was assessed for normality and the subsequent appropriate statistical test was performed as indicated in the legend of each figure.

## Results

### Inappropriate response to tissue damage in the skin during ageing

Old (≥65 years) and young (<40 years) individuals were recruited for this study, donor characteristics can be found in Table 1. Although there were more females in our older adult group, there was no difference in response between male and female older adults (Supplementary Figure 1A). Volunteers were injected intradermally with VZV skin test antigen in one arm (for the clinical score) and with saline (0.9% NaCl) in the other arm as a control. 5mm biopsies were collected from the injection sites 6 hours post-injection and compared to biopsies of normal (unmanipulated) skin (Figure 1A). We showed previously that mononuclear phagocyte numbers were increased at the site of saline injection in old but not young subjects and that this accumulation was negatively correlated with the magnitude of antigen-specific responses induced in the contralateral arm of the same individual (Vukmanovic-Stejic et al., 2018). Multiparametric flow cytometry on disaggregated skin biopsies from old donors showed no change in the number of CD11c^+^ dendritic cells in response to saline challenge (data not shown), suggesting the increase in mononuclear phagocytes was due to accumulation of monocytes.

To enumerate the changes in monocyte numbers following saline injection we assessed paired samples and expressed data as delta change from normal skin. This was to mitigate against inter-personal variation in the magnitude of responses observed (Supplementary Figure 2A). CD14 is expressed by the majority of peripheral blood monocytes (classical and intermediate). There was a significant increase in CD14^+^ cells in saline injected skin at 6 hours post-injection (Figure 1B and 1C). We next investigated the expression of CCR2, a chemokine receptor for CCL2, which is highly expressed by CD14^+^ monocytes (Patel et al., 2017; Ziegler-Heitbrock et al., 2010). There was a significant increase in number of CD14^+^CCR2^+^ cells in saline injected skin as compared to normal skin in old but not young subjects (Figure 1C). This response to saline was transient as by 24 hours post-injection there is was no significant difference in frequency of monocytes between normal and saline-injected skin (Supplementary Figure 2C). Multiparametric flow cytometry analysis of normal and saline biopsies, using our previously published panel to identify mononuclear phagocytes (Vukmanovic-Stejic et al., 2018), confirmed that there was a significant increase in CD14^+^ and CCR2^+^CD14^+^ cells in old saline-injected skin (Figure 1D).

**Figure 2:**
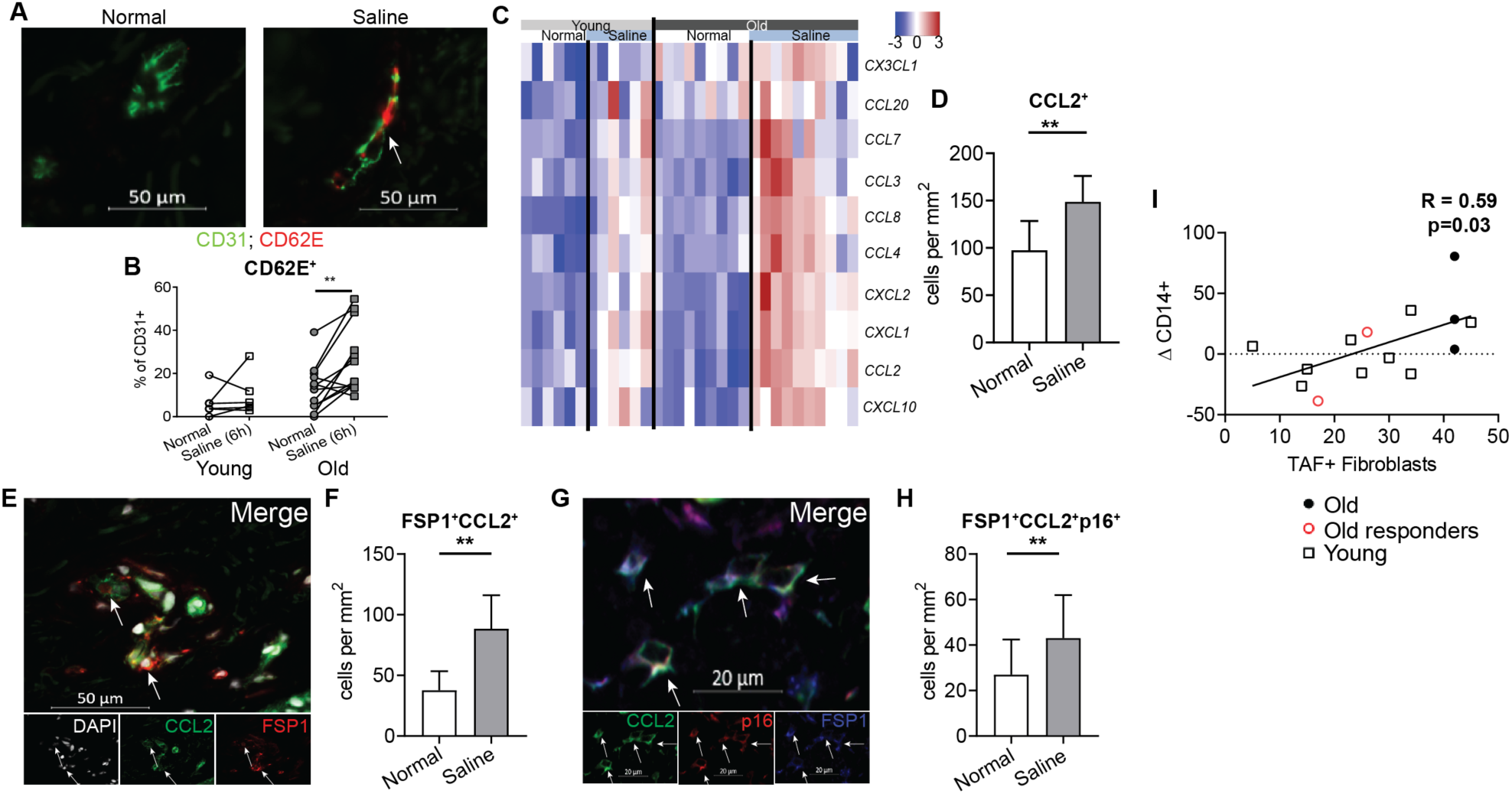
Monocyte chemoattractant expression after saline injection in the skin. **A**, representative image of normal and saline injected skin stained with CD31 (green) and CD62E (red) and **B**, cumulative data showing the frequency of CD31^+^ endothelial loops expressing CD62E. **C**, heat map showing expression of monocyte chemokines in normal then saline injected skin of young and old donors and **D**, cumulative data showing number of CCL2^+^ cells (n=10) and **E**, representative image, arrows indicate double positive cells and **F**, cumulative data showing number of FSP1^+^CCL2^+^ cells in normal and saline injected old skin (n=10). **G**, representative image showing CCL2 (green), p16 (red) and FSP1 (blue) arrows indicate FSP1^+^CCL2^+^p16^+^ cells and **H**, cumulative data of the number of FSP1^+^CCL2^+^p16^+^ cells (n=10). **I**, correlation between the number of senescent fibroblast in normal skin (defined as being Telomere associated DNA damage foci [TAF] positive;) and the change in CD14^*+*^ monocytes after saline injection **B, D, F and H**, assessed with a paired t-test. *= p<0.05, ** = p<0.01 **I**, assessed by a Spearmans rank correlation.

To investigate whether this inflammatory response after saline injection in older humans was due to sodium chloride itself or a response to needle injury, we performed intradermal injections with the same volume of air instead of saline (20μl). Air injection also induced a significant increase in the number of monocytes at the site of air challenge in old but not young subjects compared to normal skin in both groups (Supplementary Figure 2B). The accumulation of monocytes was also observed at six hours at the site of VZV injection in old but not young adults (Figure 1E). Taken together these data suggest that old subjects have a heightened inflammatory response to mild skin injury that is not observed in young individuals.

We previously showed that >85% of older individuals have a low VZV response, clinical score between 0-3, while >85% of young individuals have a score of between 3-10 (Agius et al., 2009; Vukmanovic-Stejic et al., 2015). However, a small proportion of older people (∼15%) show good cutaneous VZV responsiveness (score 3-10), a group we designated as ‘old responders’. To assess whether the CD14^+^ monocyte infiltration was a characteristic of all aged individuals or only those with reduced VZV-specific immunity, we investigated the infiltration of these cells after saline in old VZV responders (red circles) and low responders (filled circles). Old responders to VZV exhibited a similar low extent of CD14^+^ monocyte infiltration to younger adults (black squares). However, the majority of older donors with a low clinical score (filled circles) had elevated infiltration of these cells. When all the data was combined there was a significant negative correlation between the fold change in CD14^+^ monocytes in response to saline challenge and the VZV clinical score (R=-0.38 and p=0.04; Figure 1E). Collectively this indicates that the early recruitment of monocytes to the site of skin challenge in old people was associated with decreased antigen-specific immunity.

### Senescent stromal cells contribute to recruitment of monocytes in the old

We hypothesised that the increase in CD14^+^ cells in the skin was due to recruitment of circulating monocytes, since CCR2^+^ expression supported the possibility of recruitment in response to CCL2. The alternative explanation that proliferation of local tissue-resident macrophages (identified as CD163^+^) was not observed (data not shown). This is in line with murine studies showing dermal macrophages are constantly replenished from the monocyte pool (Tamoutounour et al., 2013). We first investigated the activation status of the dermal endothelium to determine whether it was permissive for monocyte recruitment. CD62E, also known as E-selectin, was used as a marker of activated endothelium as it binds to sialylated carbohydrates expressed on leukocytes and facilitates their extravasation into the tissue. Saline-injected skin of old but not young adults exhibited increased CD62E expression on the CD31^+^ endothelium when compared to normal skin (Figure 2A and B). Furthermore, transcriptional analysis of skin biopsies indicated that there was increased expression of a range of genes encoding monocyte chemoattractants including *CCL2, CXCL10, CXCL2* and *CXCL1* in old but not young saline-injected skin as compared to normal skin (Figure 2C). The increase of *CCL2* expression (one of the main monocyte chemoattractants) was confirmed by immunofluorescence staining of skin sections (Figure 2D).

We next examined the source of monocyte chemoattractants and found that resident mononuclear phagocytes were not the main source of CCL2 (data not shown). Using FSP-1 as a marker of fibroblasts, we found that there was a significant increase in the number of fibroblasts expressing CCL2 in the skin of old subjects after saline injection as compared to normal un-injected skin from old donors (Figure 2E and F). It is known that fibroblasts can produce CCL2 (Pereira et al., 2019). We and others have shown that there are increased numbers of senescent fibroblasts in the skin of older humans (Dimri et al., 1995; Pereira et al., 2019; Ressler et al., 2006) and CCL2 is a known component of the senescence associated secretory phenotype (SASP) (Campisi, 2013). Indeed, using p16 as a marker of senescence we found that FSP1^+^p16^+^ fibroblasts were the main source of CCL2 after saline injection of old donors (Figure 2G and H). Senescent fibroblasts made up 61.8% of CCL2^+^FSP1^+^ cells in normal skin which increased to 70.7% in saline-injected skin (data not shown). Although some non-senescent fibroblast evidently expressed CCL2, our previous data showed that senescent fibroblasts produce significantly more CCL2 than the non-senescent fibroblasts (Pereira et al., 2019). Therefore, the early production of SASP-related mediators by senescent fibroblasts after saline injection (including CCL2) may lead to the recruitment of inflammatory monocytes to the site of skin challenge. This interaction between senescent cells and monocyte recruitment is supported indirectly by the significant correlation between the number of senescent fibroblasts present in the skin and the extent of CD14^+^ monocyte infiltration after saline injection (Fig. 2I).

### Monocytes inhibit T_RM_ proliferation through production of Prostaglandin E2

We next investigated the mechanisms by which the recruitment of monocytes reduced antigen-specific immunity in old subjects. We first assessed the expression of inhibitory pathways in normal and saline injected skin pre- and post-Losmapimod by RNAseq. We found that saline injection (in old individuals) resulted in increased expression of a number of immunoglobulin-like transcript (ILT) receptors such as *LILRB3* (ILT5), *LILRB2* (ILT4), *LILRA2* (ILT1), *LILRB4* (ILT3) and *LILRB1* (ILT2), inhibitory ligands such as *CD274* (programme-death ligand 1; PDL-1) and *PDCD1LG2* (PDL-2), ligand for the T cell receptor TIGIT and *PVR*, as well as inducible COX2 enzyme *PTGS2* (Figure 3A).

**Figure 3:**
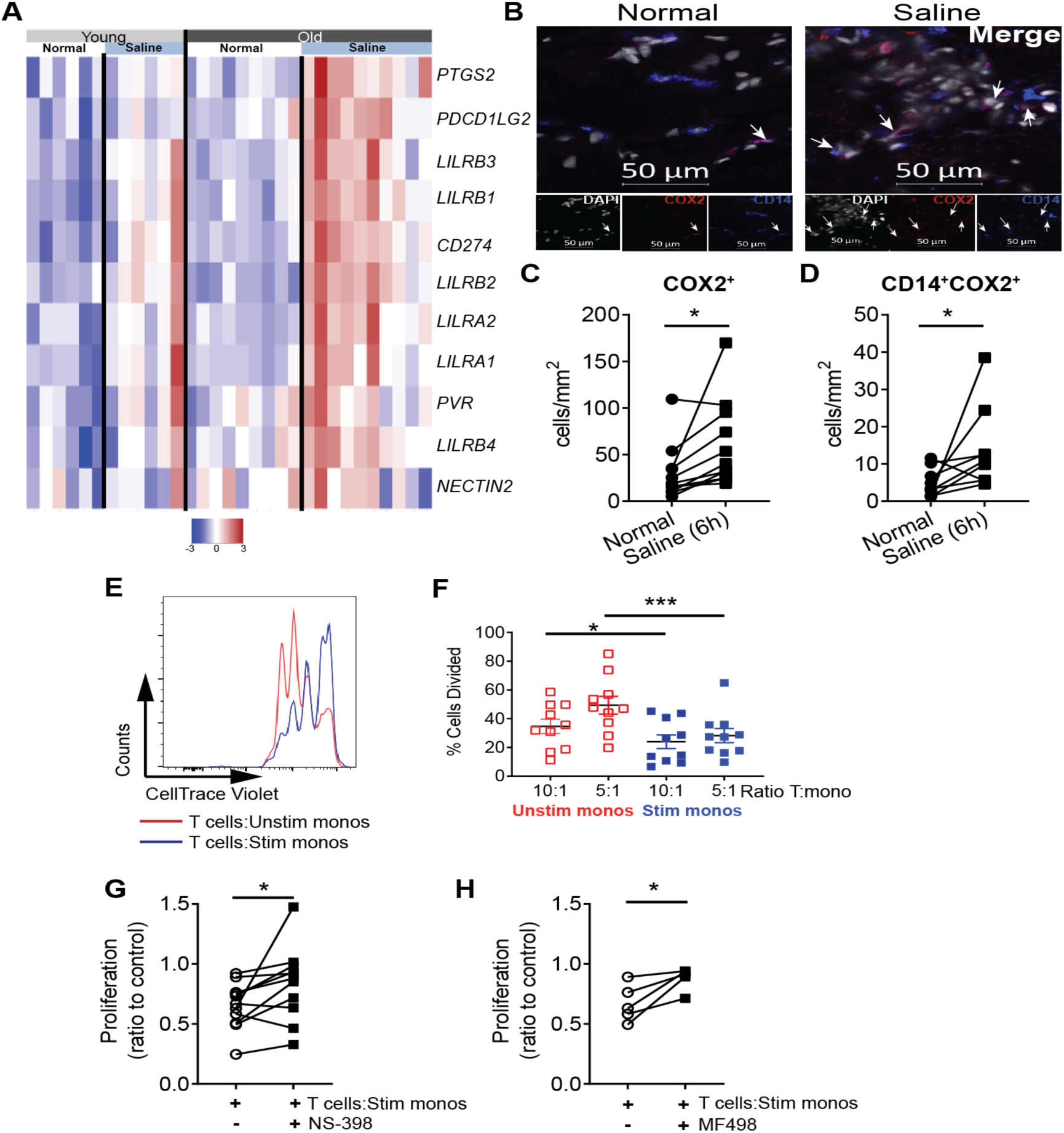
CD14^+^COX2^+^ monocytes inhibit T cell proliferation via PGE_2_. **A**, Heatmap showing the expression of genes associated with inhibitory mechanisms in normal and saline injected of old and young donors **B**, representative staining of DAPI (white), COX2 (red) and CD14 (blue) in normal and saline injected skin and **C**, cumulative data of total COX2 expressing cells and **D**, number of CD14^+^COX2^+^ cells in normal and saline injected skin. **E**, Monocytes were negatively isolated from the peripheral blood and cultured with and without LPS for 3 hours then subsequently co-cultured with autologous T cells (pre-labelled with CellTrace Violet) that were activated with plate-bound CD3 and IL-2, proliferation was assessed at day 4. A representative flow plot of CellTrace violet dilution in CD4^+^ T cells co-cultured with unstimulated and stimulated monocytes is shown in **E** and **F**, cumulative data showing percent of cells divided. A similar co-culture experiment was performed in the presence or absence of the COX2 inhibitor NS-398 or **G**, the EP4 receptor inhibitor MF498 **H**. In **C**,**D and F**, analysed with a paired t-test. **G** and **H** analysed with Wilcoxon matched-pairs signed rank test * = p<0.05; *** = p<0.001

*PTGS2* (encoding COX2) was one of the most highly upregulated genes in saline-injected skin of the old subjects (Vukmanovic-Stejic et al., 2018) and elevated COX2 expression following saline injection was confirmed at the protein level by immunohistology (Figure 3B and C). When staining for COX2 was performed in combination with CD14 there was a significant increase in CD14^+^COX2^+^ cells in saline injected as compared to normal skin (Figures 3D).

To demonstrate that COX2 expressing monocytes can inhibit the activation of CD4^+^ T cells, we first induced COX2 expression in these cells that were isolated from peripheral blood by stimulation with LPS (1ng/ml) for 3 hours. LPS treatment significantly increased COX2 expression in monocytes compared to untreated controls (Supplementary Figure 3) (Dean et al., 1999; Mestre et al., 2001). Unstimulated and LPS-stimulated monocytes were co-cultured with autologous peripheral CD4^+^ T cells in the presence of anti-CD3 and IL-2 and proliferation was assessed at after four days. Stimulated monocytes inhibited CD4^+^ T cell proliferation as compared to control (unstimulated) monocytes (Figure 3E-F). However, production of the cytokines IFNy, IL-2 and IL-10 was not affected at day 4 post-culture (data not shown). To determine if COX2 expression was involved in the inhibition of CD4^+^ T cell proliferation, a COX2-specific inhibitor NS-398 was added to the co-cultures. Treatment with NS-398 led to a significant increase in the proliferation of the CD4^+^ T cells (Figure 3G). COX2 is involved in the production of a range of lipid mediators, the most common being PGE_2_ (Kalinski, 2012). To establish if PGE_2_ was involved in the inhibition of CD4^+^ T cell proliferation, we blocked the receptor for PGE_2_, EP4, on CD4^+^ T cells using the inhibitor MF-498. The blockade of PGE_2_ signalling significantly increased the proliferation of the CD4^+^ T cells in the presence of inflammatory monocytes (Figure 3H). To confirm that PGE_2_ did not alter the monocyte phenotype we cultured monocytes with PGE_2_ and assessed their inhibitory receptor (CD112, CD155, Galectin 9, ILT3 and PDL-1, PDL-2) expression and cytokine production. We did not observe any significant effects on the expression of these receptor expression in monocytes (data not shown), confirming the effect of PGE_2_ was on the CD4^+^ T cells directly. To assess if the above observations were in operation in the skin of old donors *in situ*, we investigated EP4 expression by skin-resident CD4^+^ T_RM_ cells in older volunteers. Immunofluorescence staining was performed on normal skin to determine expression of EP4 on CD4^+^ T_RM_ (as identified by co-expressing CD69; Figure 4A). The majority of CD4^+^ T_RM_ expressed EP4 (81.2%), indicating that they have the capacity to respond to PGE_2_ produced by monocytes. We next assessed the co-localisation of CD14^+^ monocytes and CD4^+^T_RM_ cells in normal and saline injected skin from old subjects. In normal skin, CD14^+^ monocytes were on average 15.86μm away from CD4^+^ T_RM_, however after saline injection the monocytes were significantly closer to the CD4^+^ T_RM_ (9.47μm; Figure 4B and C). This increased proximity was likely due to increase in the number of monocytes in the skin after saline injection. To confirm that monocytes have the ability to suppress skin CD4^+^ T_RM_ proliferation, CD4^+^ T_RM_ were isolated from the skin using suction blister technology as described previously (Figure 4D) (Akbar et al., 2013). Skin CD4^+^ T_RM_ cells were pre-labelled with CellTrace Violet and stimulated with CD3 and IL-2 in the presence of either stimulated or unstimulated monocytes for four days. We observed a significant reduction in CD4^+^ T_RM_ cell proliferation in the presence of stimulated when compared to unstimulated monocytes (Figure 4E and F) supporting the hypothesis that COX2^+^ monocytes have the capability to suppress T_RM_ proliferation.

**Figure 4:**
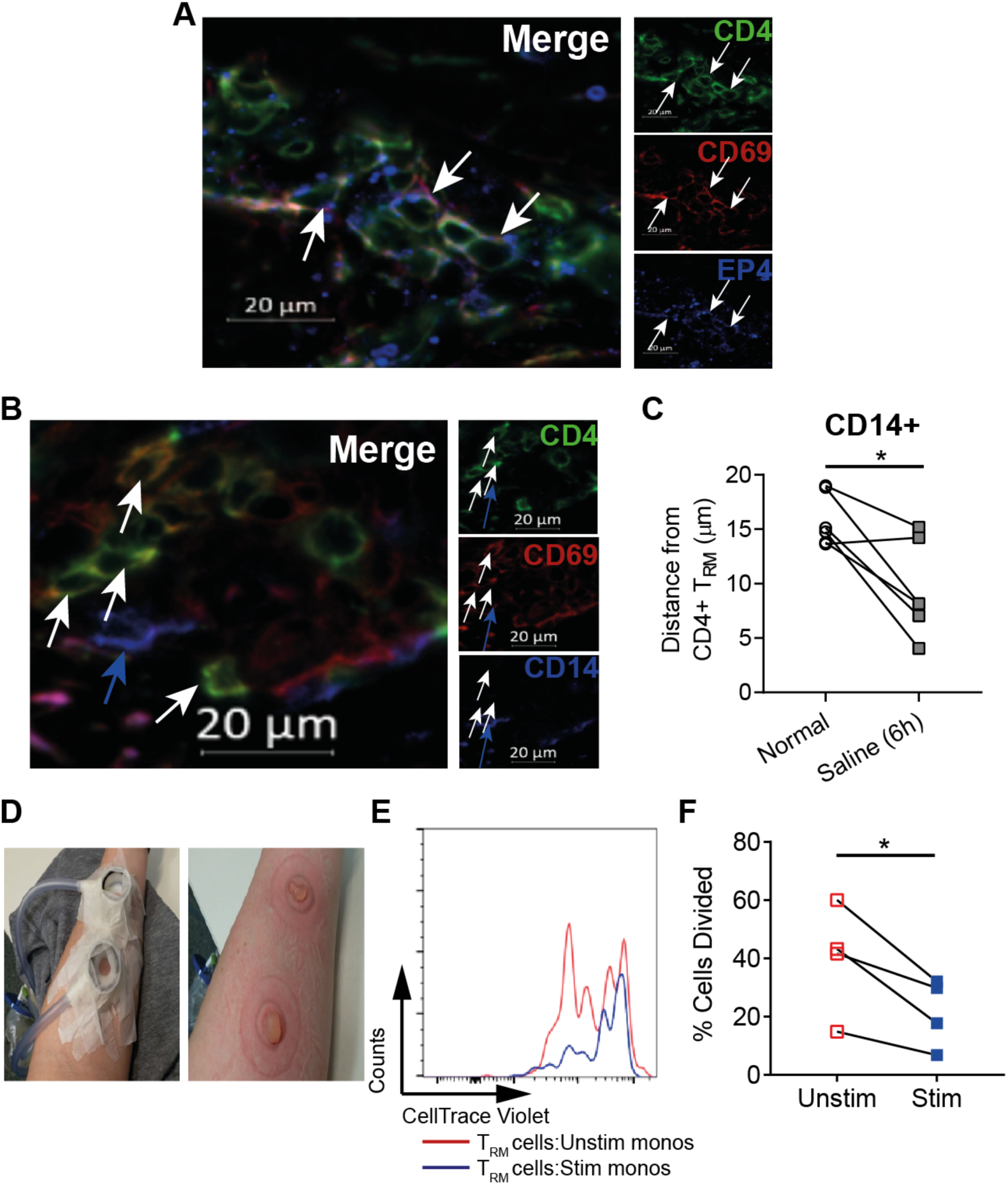
PGE2 production by monocytes inhibits skin CD4^+^ T_RM_ cell proliferation. **A**, representative staining of CD4 (green), CD69 (red) and EP4 (blue) in normal skin, white arrows indicate CD4^+^ T_RM_ cells which co-express EP4. **B**, representative staining showing co-localization of CD4^+^ T_RM_ cells and CD14^+^ monocytes (blue) and **C**, cumulative data showing distance of CD14^+^ monocytes from T_RM_ in normal and saline injected skin from older adults. Monocytes were negatively isolated from the peripheral blood and cultured with and without LPS for 3 hours then subsequently co-cultured with skin T_RM_ cells (pre-labelled with CellTrace Violet), which were collected from suction blisters (representative image in **D**) then activated with plate-bound CD3 and IL-2 and proliferation was assessed at day 4. **E**, representative flow plot of CellTrace violet dilution in T_RM_ cells co-cultured with unstimulated (red line) or stimulated (blue line) monocytes. **F**, Cumulative data on CD4^+^ T_RM_ proliferation in the presence of stimulated or unstimulated monocytes, assessed at day 4. **C**, and **F**, were analysed with a paired t-test. * = p<0.05

### p38 MAP kinase inhibition significantly reduces CCL2 production from senescent fibroblasts

We showed previously that treatment of old subjects with the anti-inflammatory p38-MAPK inhibitor, Losmapimod, significantly enhanced the response to VZV antigens in the skin (Vukmanovic-Stejic et al., 2018). To investigate if p38-MAPK inhibition altered CCL2 production and inflammatory monocyte recruitment in the skin, RNAseq analysis was performed on biopsies collected from normal skin and from skin 6 hours after saline injection with saline both before and after Losmapimod treatment (study design Figure 5A). We stratified our subjects in the RNAseq analysis according to the extent of clinical score improvement. The individuals that had no improvement in clinical score are represented as white and those who had the biggest improvement in clinical score after Losmapimod treatment represented by dark green (Figure 5B). A list of all differentially expressed genes can be found in Supplementary Table 1. Saline injection resulted in a large pro-inflammatory response with a significant increase in the expression of inflammatory genes such as *CCL2, CCL8, FOSL1, IL1B* and *TREM1* at 6 hours pre-Losmapimod treatment (Figure 5C). This response to saline was reduced after Losmapimod pre-treatment (Figure 5C) and the top six upregulated genes that were significantly different between normal and saline injected skin were no longer significantly different post-Losmapimod treatment (Supplementary Figure 4A). In addition, RNAseq analysis showed that the expression of monocyte chemoattractants (induced in response to saline injection) was reduced post-Losmapimod treatment (Supplementary Figure 4B). Furthermore, after Losmapimod there was a significant reduction in the number of CCL2^+^ cells and the number of CCL2^+^ senescent fibroblasts as defined as being p16^+^FSP-1^+^ (Figures 5D, 5E). We confirmed experimentally that the treatment of senescent dermal fibroblasts with Losmapimod *in vitro* significantly reduced their CCL2 production, suggesting Losmapimod could have a direct effect on senescent fibroblast *in vivo* (Supplementary Figure 4C). As a result of the reduced CCL2 there was significantly reduced monocyte infiltration in saline injected skin Post-Losmapimod (Figure 5F). In addition, when VZV injected skin (6 hour post-injection) was assessed there was a significant reduction in the number of monocytes after Losmapimod treatment. This data indicates that p38-MAPK inhibition significantly reduces CCL2 production from senescent fibroblasts reducing the accumulation of monocytes at sites of saline or VZV antigen challenge in the skin (Figure 5G).

**Figure 5:**
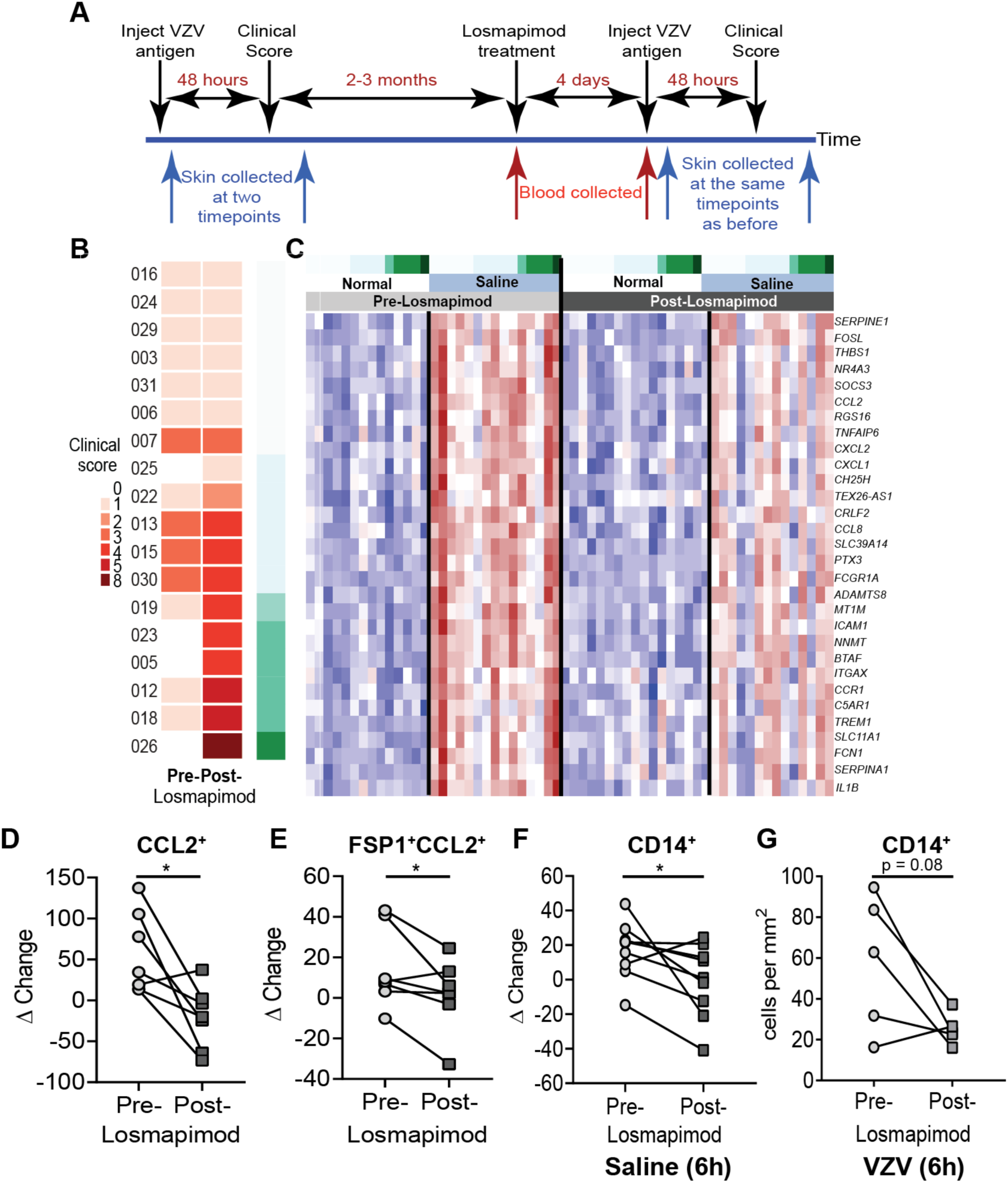
Reduced inflammatory monocyte recruitment to the skin by Losmapimod pre-treatment. **A**, Losmapimod clinical study diagram. Black arrows indicate study visits, red arrows indicate blood samples collected and blue arrows indicate skin biopsies taken. **B**, Colour coding of the clinical response of each individual after VZV challenge both before and after Losmapimod pre-treatment (pink/brown). Further colour coding was introduced to identify the extent of improvement where dark green identifies the biggest improvement and white shows no improvement in the skin. **C**, This coding was used in conjunction with the transcriptomic signatures in normal and saline injected skin before and after Losmapimod treatment. The top 30 genes upregulated in saline injected skin as compared to normal skin before pre-Losmapimod. **D**, Fold change of CCL2^+^ cells and **E**, FSP1^+^CCL2^+^ cells in saline injected pre- and post-Losmapimod treatment. **F**, fold change CD14^+^ cell in the skin in response to saline injection and **G**, number of CD14^+^ monocytes in VZV injected skin (6 hours after injection) pre- and post-Losmapimod treatment. Significance in **D-G** assessed using a paired t-test. * = p<0.05

### p38 MAP kinase inhibition significantly inhibits PGE_2_ that restores T cell proliferation

COX2 expression has been shown to be downstream of p38-MAPK signalling (Dean et al., 1999; Guan et al., 1998), implying that Losmapimod may act by inhibiting COX2 expression *in vivo*. To test this, we performed Western blot analysis of LPS activated monocytes in the presence or absence of Losmapimod. Losmapimod significantly decreased the LPS induced expression of phopho-p38 MAPK and COX2 *in vitro* (Figures 6A and B). In addition, PGE_2_ production was also significantly decreased after Losmapimod treatment (Figure 6C). Furthermore, the addition of Losmapimod to co-cultures of CD4^+^ T cells that were activated in the presence of LPS treated monocytes (Figure 6D) enhanced their proliferation to a similar extent to that observed after direct COX2 or EP4 inhibition (Figure 3G and H). In line with these observations, the increase in CD14^+^COX2^+^ cells after saline injection was also inhibited after post-Losmapimod treatment *in vivo* (Figure 6E). These data collectively suggest that *in vivo* Losmapimod can restore CD4^+^ T cell responses in part through the inhibition of COX2 expression.

**Figure 6:**
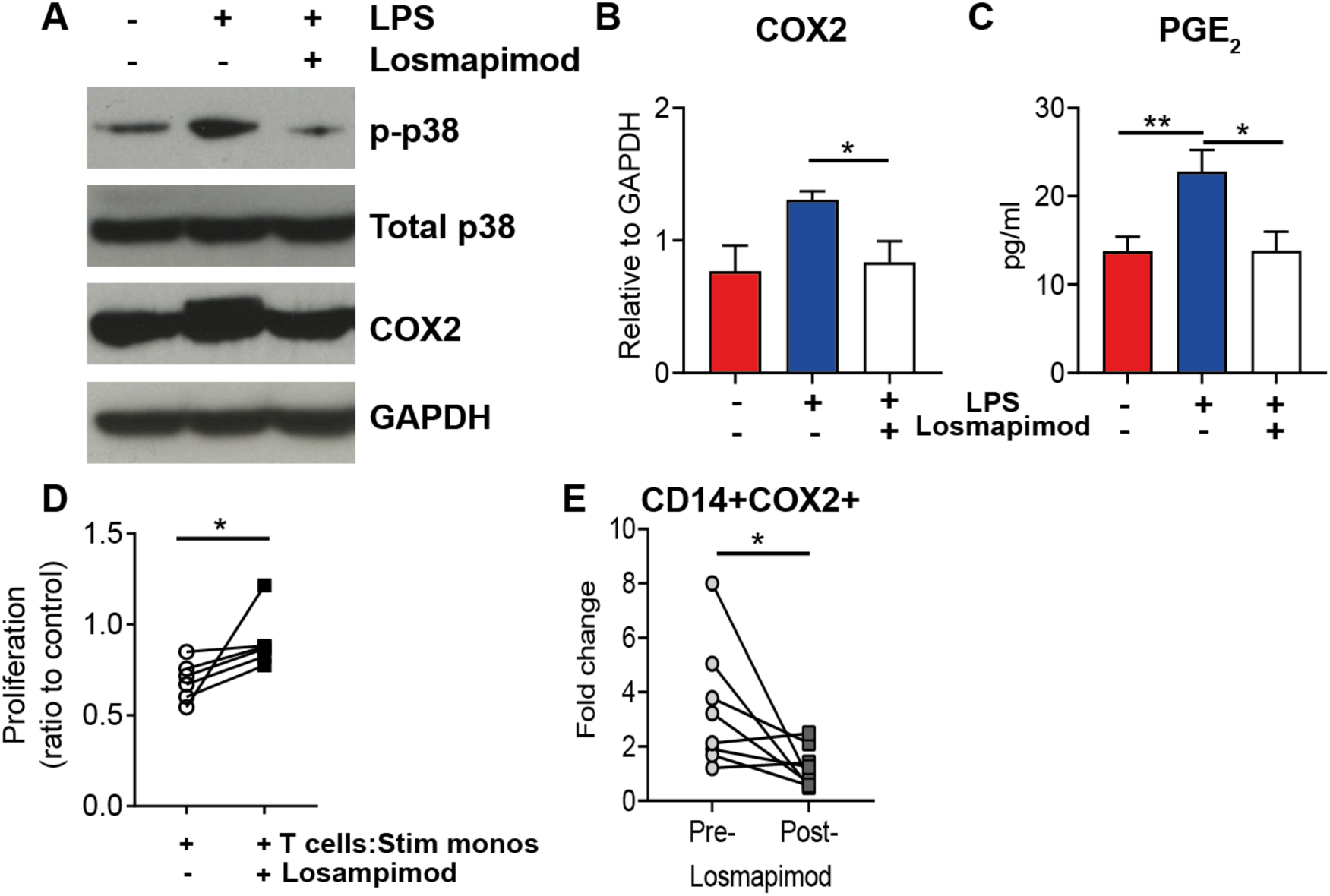
Losmapimod restores T cell function by inhibiting monocyte COX2 and PGE_2_ production. Monocytes were negatively isolated from the peripheral blood and cultured with and without LPS and in the presence or absence of Losmapimod for 3 hours, pellets were collected and western blot performed **A**, representative blot of phospho-p38 MAP Kinase (p-p38), total p38 MAP Kinase (p38), COX2 and GAPDH. **B**, cumulative data of COX2 expression relative to GAPDH (n=3) and **C**, Prostaglandin E2 (PGE_2_) production. Monocytes were negatively isolated from the peripheral blood and stimulated with LPS (blue) and without LPS (red) or with LPS in the presence of Losmapimod (white) for 3 hours then subsequently co-cultured with autologous T cells (pre-labelled with CellTrace Violet) in the presence of plate-bound CD3 and IL-2 proliferation was assessed at day 4. **D**, cumulative data showing percent of cells divided. **E**, fold change of **i**nfiltration of CD14^+^COX2^+^ cells between normal and saline injected skin before and after Losmapimod treatment. **B, and C**, analysed by a one-way ANOVA with Tukey’s multiple comparison test and **D, and E**, analysed with a Wilcoxon matched-pairs signed rank test * = p<0.05

### p38 MAP kinase inhibition significantly increases T cell infiltration into the site of VZV challenge

We assessed the clinical score of 42 old subjects after VZV antigen challenge in the skin both before and after Losmapimod treatment. We confirmed and extended our previous observations that treatment with Losmapimod significantly increased VZV clinical score (n=42; Figure 7A). No difference was observed between male and female donors in their response to Losmapimod treatment (Supplementary Figure 1B). Losmapimod treatment also significantly decreased serum CRP in all individuals, irrespective of whether or not they increased their skin response to VZV challenge (Figure 7B).

**Figure 7:**
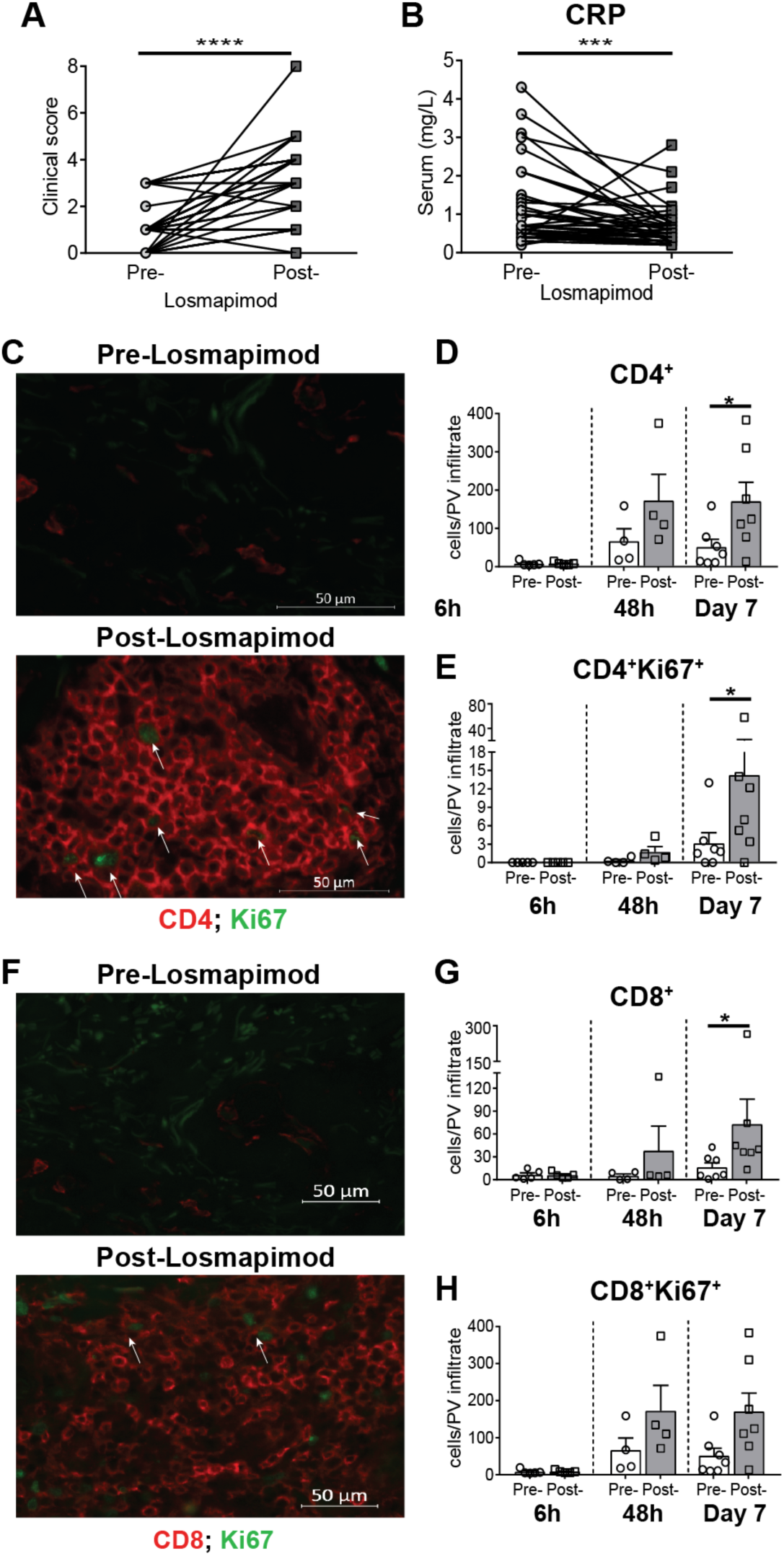
Losmapimod treatment increases VZV induced T cell responses in the skin. **A**, VZV clinical score in all donors (n=42) and **B**, serum CRP concentrations pre- and post-Losmapimod treatment. **C**, representative images of CD4 (red) and Ki67 (green) staining at day 7 days after VZV challenge with and without Losmapimod pre-treatment. **D**, cumulative data of CD4^+^ staining and **E**, CD4^+^Ki67^+^ staining at different times after VZV challenge, pre and post Losmapimod pre-treatment. **F**, representative image of CD8 (red) and Ki67 (green) staining at day 7 after VZV challenge and **G**, cumulative data of CD8^+^ staining and **H**, CD8^+^Ki67^+^ staining at different times after over the time course pre and post Losmapimod pre-treatment. **D, E, G, and H** analysed with a paired t-test. * = p<0.05

We showed previously that there is a significant association between the clinical score after VZV antigen challenge and the accumulation of T cells at the site of injection (Agius et al., 2009; Vukmanovic-Stejic et al., 2018). We found that Losmapimod pre-treatment was associated with a significant increase in CD4^+^ (Figure 7C,7D) and CD8^+^ (Figure 7F, 7G) T cell infiltration into the skin 7 days after VZV injection. The increased number of CD4^+^ and CD8^+^ T cells in perivascular infiltrates was associated with an increased proliferation of these cells *in situ* indicated by Ki67 expression (Figures 7E and H) (Vukmanovic-Stejic et al., 2015). Taken together our data show that short-term blockade of inflammation using the p38 MAP kinase inhibitor Losmapimod can significantly enhance the cutaneous immune response to antigen challenge in older adults.

## Discussion

Older humans exhibit decreased recall responses to antigens in the skin (Agius et al., 2009). Furthermore, older adults who had the greatest tendency to mount inflammatory responses after mild tissue injury induced by saline injection had the lowest response to recall response to VZV antigen challenge in the skin (Vukmanovic-Stejic et al., 2018). This suggested that increased propensity to exhibit inflammatory responses leads to inhibition of VZV-specific T cell responses in the skin. The fact that the temporary blockade of inflammation in these old individuals with a p38-MAPK inhibitor *in vivo* enhanced the response to VZV antigen challenge in the skin supports this contention (Vukmanovic-Stejic et al., 2018). However, the source of inflammation in the skin of older subjects or the mechanism by which the inflammation inhibited T_RM_ cell function was not known.

We now show that senescent fibroblasts in the skin of old subjects secrete an array of chemokines including CCL2 in response to tissue injury, such as those induced by injecting saline, VZV or air. This leads to recruitment of inflammatory monocytes into the skin that inhibit skin T_RM_ activation in part by secreting lipid mediators such as PGE_2_ via a COX2-dependant pathway. However, short-term systemic p38-MAPK blockade induced by treating old donors with Losmapimod, leads to significantly reduced CCL2 (and other chemokine) production by senescent fibroblasts, reduced monocyte infiltration and reduced COX2 and PGE2 expression in response to injection of saline or VZV antigens. This results in significant improvement of VZV-specific cutaneous T cell responses (Figure 8). Therefore, p38-MAPK inhibition can act by significantly reducing the recruitment of inflammatory monocytes into the skin and also by inhibiting the function of those that still manage to extravagate into the tissue.

**Figure 8:**
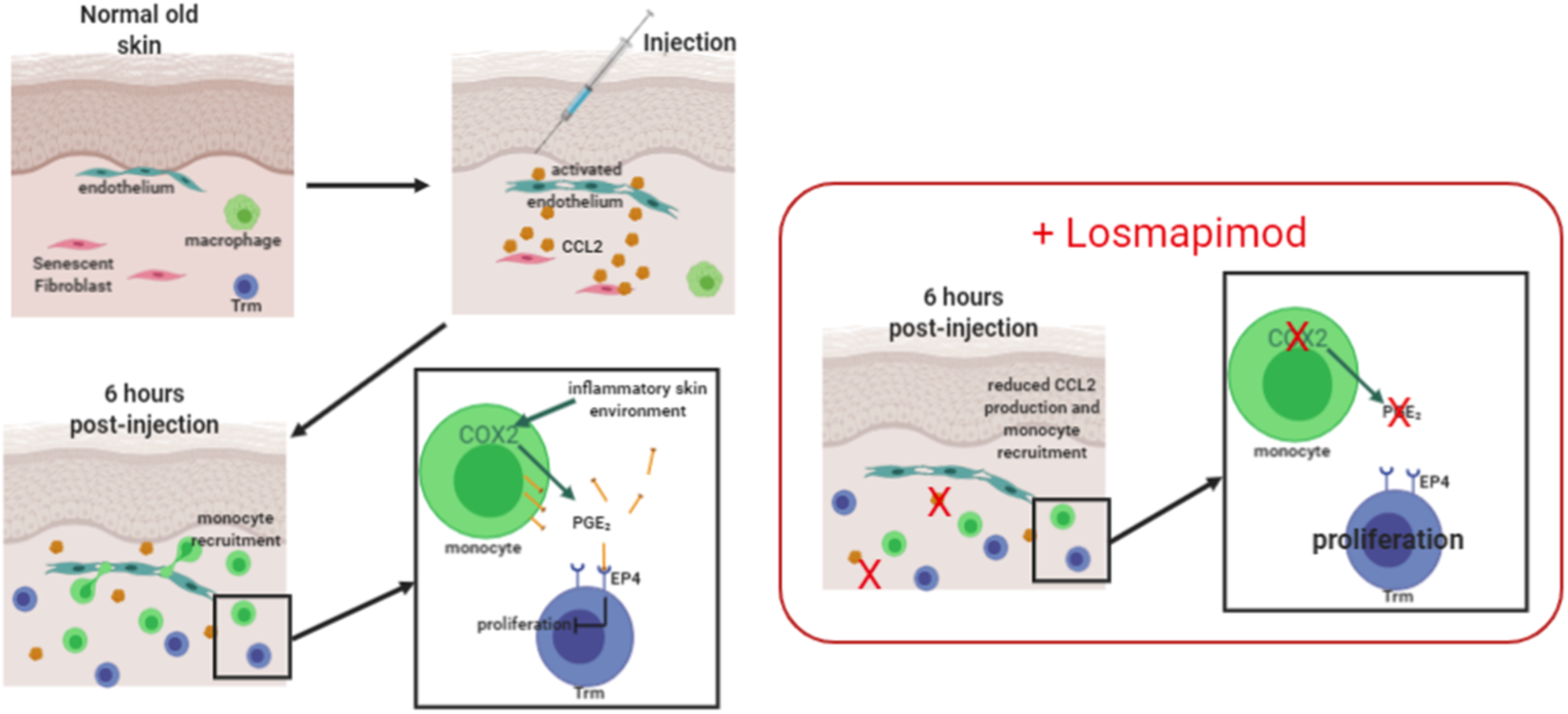
Schematic model of monocyte inhibition of cutaneous immunity in older adults. In older adults the mild damage induced by injection (of air, saline or VZV) results in production of CCL2 from senescent fibroblasts and recruitment of monocytes into the skin 6 hours post injection. The recruited monocytes inhibit T_RM_ proliferation via upregulation of COX2 and subsequent production of the lipid mediator Prostaglandin E2 (PGE_2_). The treatment of older adults with the p38 MAPK inhibitor, Losmapimod, results in reduced CCL2 production from the senescent fibroblasts and reduced monocyte infiltration. In addition, p38 MAPK is upstream of COX2 signalling, therefore the treatment with Losmapimod also inhibits COX2 upregulation and reduced PGE_2_ production. This significantly increases the T cell response to VZV in the skin of older subjects

Ageing leads to the accumulation of senescent cells that is observed in many tissues including the skin (Pereira et al., 2019; Ressler et al., 2006); therefore, the recruitment of inflammatory monocytes in response to signals from senescent fibroblasts may contribute to age-related decreased immunity in other organs as well as the skin. CCL2 is involved in the recruitment of monocytes into the skin and it is a component of the SASP (Pereira et al., 2019). The significant correlation between the number of senescent fibroblasts that were present in the skin before injection and the extent of CD14^+^ monocyte infiltration indirectly supports the interaction between these cells. Although the abundance of senescent cells increases in the skin during ageing (Dimri et al., 1995; Pereira et al., 2019; Ressler et al., 2006) there is no overt expression of inflammatory mediators in un-manipulated skin of old subjects (Vukmanovic-Stejic et al., 2015). This suggests that at the steady state, the propensity to mount inflammatory responses including the SASP is inhibited in this tissue. In support of this idea there are increased numbers of Foxp3^+^ T regulatory cells (Tregs) and increased expression of the inhibitory receptor PD-1 by cutaneous T_RM_ cells, which may contribute to the control of inflammation in this tissue I older individuals (Vukmanovic-Stejic et al., 2008; Vukmanovic-Stejic et al., 2015). One possibility is that Tregs may inhibit the secretion of inflammatory mediators by senescent tissue stromal cells and this requires further investigation. The breakdown of homeostasis induced by tissue injury, may override pre-existing inhibitory processes and allow the secretion of SASP-related factors leading to the subsequent events that we have described

We found that skin infiltrating monocytes secrete PGE_2_ via a COX2 dependent pathway. PGE_2_ has disparate roles in immune responses as it has been shown to be pro-inflammatory (causing oedema formation and pain) as well as contributing to resolution of inflammation (Kalinski, 2012). Here we show that PGE_2_ produced by COX2 expressing inflammatory monocytes inhibits T cell proliferation *in vitro* and may directly inhibit antigen-specific T_RM_ activation in the skin. PGE_2_ has been shown previously to directly inhibit CD4^+^ T cell proliferation in humans (Okano et al., 2006). Furthermore, this inhibition may occur via increased expression of COX2 and production of PGE_2_ by monocytes (Chen et al., 2011). During chronic viral infection PGE_2_ can also directly suppress CD8^+^ T cells delaying viral clearance (Chen et al., 2013). We now show that PGE_2_ can directly suppress the activation and proliferation of human skin derived T_RM_ cells. PGE_2_ may also induce the expansion and increase the function of Foxp3^+^ Tregs (Baratelli et al., 2005; Sharma et al., 2005) explaining why these cells increases in the skin during ageing (Vukmanovic-Stejic et al., 2008; Vukmanovic-Stejic et al., 2013).

Elevated PGE_2_ concentrations are also found at the sites of chronic viral infection and virus-specific T cells in the liver of Hepatitis B infected individuals express the PGE_2_ receptor EP4 (Chen et al., 2015; Okano et al., 2006). The observation that PGE_2_ is found in tumour sites prompted the suggestion that COX2 inhibitors may be useful in the treatment of colon cancer and basal and squamous cell carcinoma (Pandeya et al., 2019; Steinbach et al., 2000). In addition, PGE_2_ may promote immunosuppression by inducing the anti-inflammatory cytokine IL-10 by mononuclear phagocytes (MacKenzie et al., 2013; Nakanishi and Rosenberg, 2013). We now show that the PGE_2_ receptor, EP4, is expressed by the majority of skin-resident T_RM_ that facilitates the inhibition of these cells by this mediator.

Monocytes were historically regarded as precursors of macrophages, however more recent data shows that they can be effector cells in their own right (Cros et al., 2010; Goritzka et al., 2015). We confirmed previous studies showing that monocytes can inhibit antigen-specific responses (Sammicheli et al., 2016; Watanabe et al., 2017). Monocytes were shown to be increased in the leukocyte infiltrates in the lungs of patients with severe COVID-19 symptoms (Minfeng Liao and Li, 2020). Since the majority of the patients with most severe symptoms are older (Stephanie Bialek, 2020), it will be important to determine the mechanism by which these “inflammatory” monocytes are recruited into the lung. It is possible that once inflammatory monocytes are recruited into the lungs of these patients they may inhibit the function of virus-specific T cells, by the mechanism that we have described, thus preventing appropriate anti-viral immunity (Stephanie Bialek, 2020). Relevant to our current observations, it has been shown that senescent cells increase in the lungs with age-dependent pathology (Barnes et al., 2019) and it will be important to understand if these cells are involved with the recruitment of these inflammatory monocytes in patients with severe COVID-19 infection. Although CCR2^+^ monocytes can contribute to the inflammatory environment in the skin, they are also actively involved in the resolution of inflammation, for example in in the gut (Farro et al., 2017). Therefore, microenvironmental cues may determine the function of monocytes that are recruited into different tissues.

Excessive inflammation has been shown to inhibit antigen-specific immunity (Pence et al., 2012) and reduce vaccination efficacy (Fourati et al., 2016; Muyanja et al., 2014; Parmigiani et al., 2013). We have shown that the effects of inflammation on inhibiting immunity can be reversed with a p38-MAPK inhibitor. Studies with other inhibitors of inflammation such as mTOR inhibitor, rapamycin have shown that they can enhance the response to influenza vaccination in older subjects by 20% (Mannick et al., 2014). Additionally, in non-human ageing models, metformin and rapamycin can increase life-span and immune function (Cabreiro et al., 2013; Miller et al., 2011; Neff et al., 2013). In addition, in a murine model of breast cancer COX2 inhibitor could improve DC-based vaccine efficacy by increasing IFNγ production and cytotoxic CD8 function (Basu et al., 2006). Therefore, many anti-inflammatory drugs are already licenced for use in humans or have been through phase III clinical trials. Our observations with the p38-MAPK inhibitor Losmapimod provide a proof-of-principle demonstration that the efficacy of these drugs could be tested for their ability to prevent the deleterious effects of inflammageing resulting in the enhancement of antigen-specific immunity in older humans *in vivo*.

## Author contribution

**ESC** designed, performed experiments and wrote the manuscript. **MVS** was involved in the overall design of the study, experimental design and wrote the manuscript. **HT** and **OD** performed the experiments. **BBS, NM** and **TCF** performed the bioinformatic analysis of the RNA-seq samples. **PS** and **JG** performed clinical procedures and sample collection. **MHR** was the clinical lead for the study and was involved with scientific discussions. **DG** advised experimental design for COX2 and PGE_2_ work. **ANA** was involved in the overall design of the study, initiated and coordinated the collaborative interaction between the different research groups, interpreted the data, contributed writing and edited the manuscript.

We thank Dr. Iain Laws, Dr. Ruchira Glaser, Dr Lea Sarov-Blat and Dr. Robert Henderson at GSK, Dr Veronique Birault at the Francis Crick Institute and Professor Mahdad Noursadeghi at University College London for support in developing this project. We are also grateful to Glaxo-Smith Kline for providing the drug Losmapimod (MICA agreement with the MRC) and to the Losmapimod team at GSK for organizing the dispatch of the clinical supply of the drug in this Investigator Led study. We would especially like to thank the blood and skin donors who volunteered for this study and to our research nurses Ms Megan Harries-Nee and Ms. Michelle Berkley for their outstanding work

## Abbreviations

CBA: cytometric bead array
CRP: C reactive protein
COX2: Cyclooxygenase 2
DC: Dendritic Cell
LPS: Lipopolysaccharide
ILT: Immunoglobulin-like transcript
p38-MAPK: p38 mitogen-activated protein kinase
PD-L: Programme-death ligand
PGE_2_: Prostaglandin E2
PV: perivascular infiltrate
SASP: senescence associated secretory phenotype
Tregs: T regulatory cells
T_RM_: T resident-memory cells
VZV: Varicella Zoster Virus

