## Supplementary figures and tables for "Monocyte-derived Prostaglandin E2 inhibits antigen-specific cutaneous immunity during ageing"

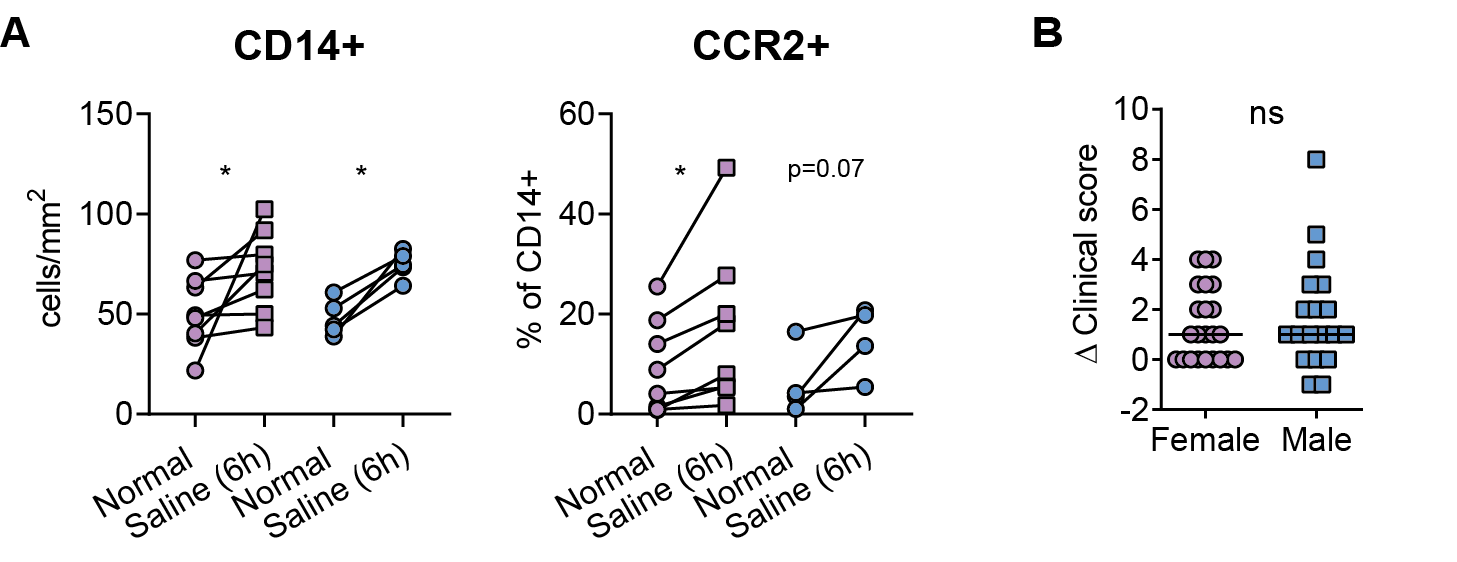

**Supplementary Figure 1: No gender difference observed in older adults in response to saline or Losmapimod**

**A,** cumulative data showing the number of CD14^+^ and frequency of CCR2^+^ CD14^+^ cells in females (pink) and males (blue). **B,** Change in clinical score after Losmapimod treatment in female (pink) and male (blue), line indicates median. Data assessed by paired t-test within donor and non-paired t test for comparison between female and males * = p<0.05.

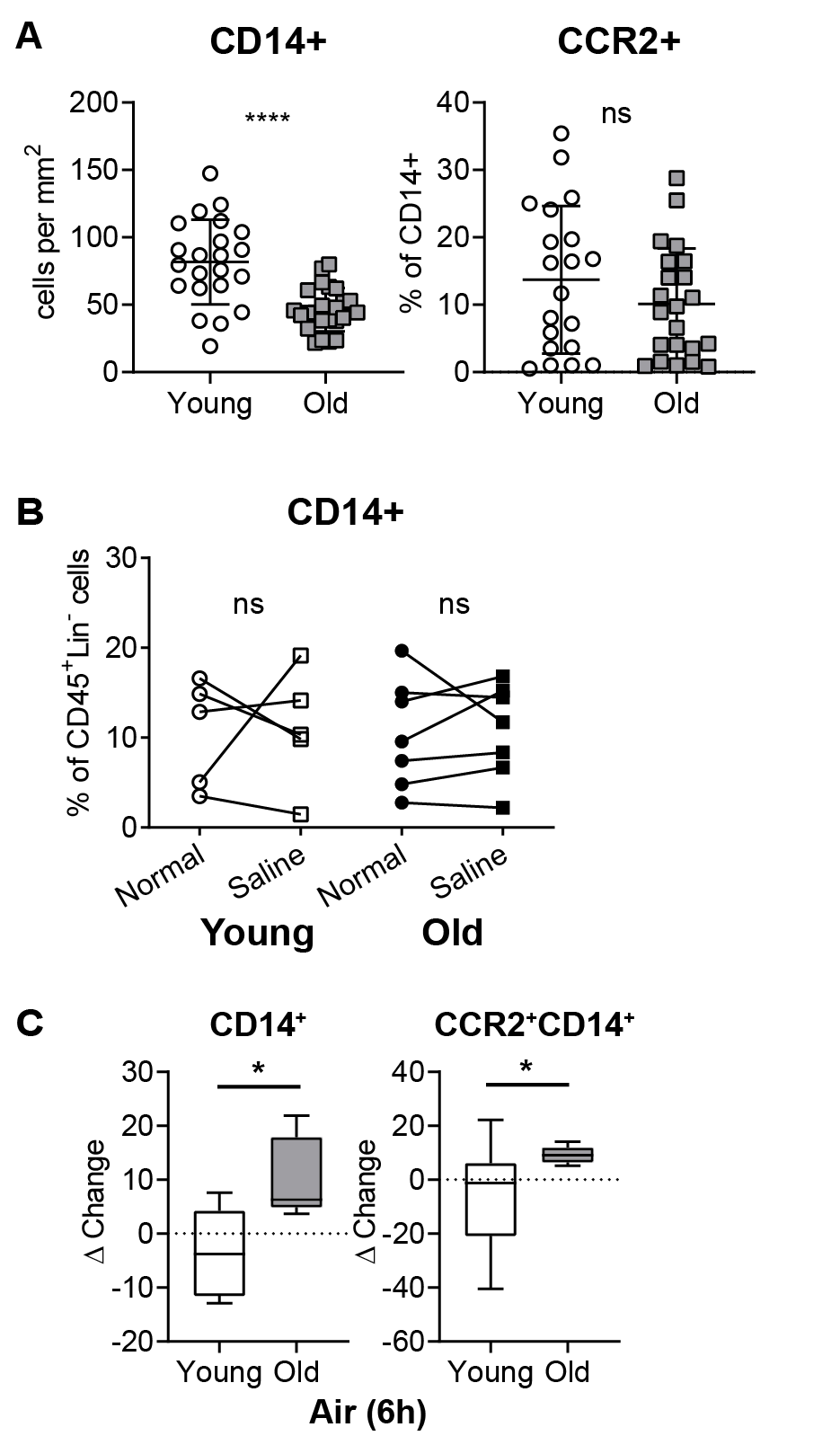

**Supplementary Figure 2: Baseline and 24 hours post-saline injection monocyte populations**

**A,** Comparison of baseline monocyte populations in the skin of based on CD14^+^ and CCR2^+^ expression in young and old individuals . **B,** Normal and Saline-injected skin were collected 24 hours post-injection from 5 young (average age 30 years; 3 male and 2 females) and 7 old (average age 70 years; 2 males and 5 females). Samples were digested overnight and assessed the following day by multiparametric flow cytometry. Cells that were CD45+HLA-DR+and CD3, CD56, CD19 and CD20 negative were considered to be mononuclear phagocytes, within this population CD14+ monocytes were assessed. **C,** Cumulative delta change in number of CD14+ or CCR2+CD14+ cells in air-injected skin from young and old volunteers as compared to normal skin. Data assessed by paired t-test * = p<0.05; **** = p<0.0001.

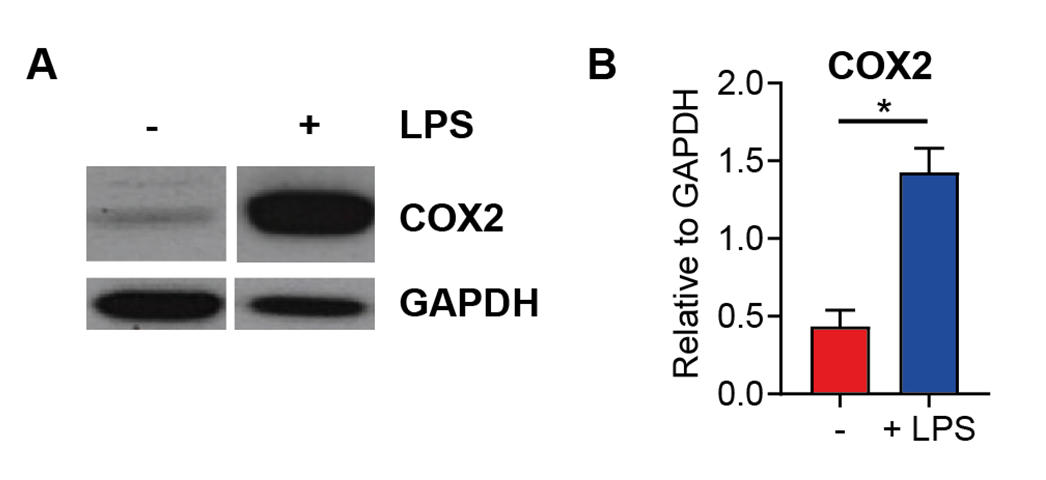

**Supplementary Figure 3: LPS treatment increases COX2 expression**

Monocytes were negatively isolated from the peripheral blood and cultured with and without LPS for 3 hours then cell pellets were collected and western blot performed **A**, representative image and **B,** cumulative data of COX2 expression relative to GAPDH (n=7). **B,** analysed with a Wilcoxon matched-pairs signed rank test * = p<0.05

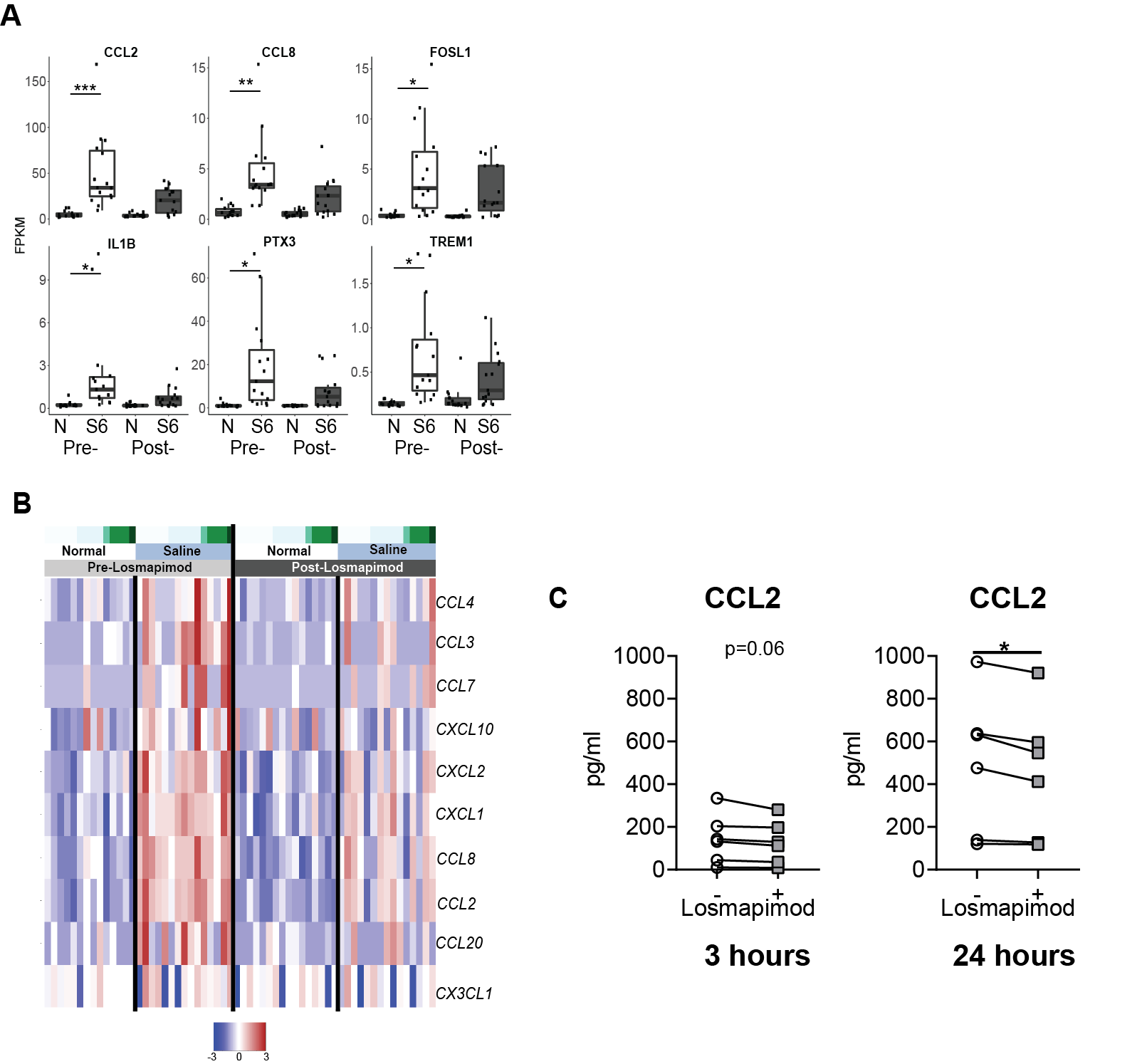

**Supplementary Figure 4: Losmapimod reduces the expression of monocyte chemokines**

**A,** top six genes upregulated in saline injected as compared to normal skin, when ranking by average fold change for genes significantly upregulated (adj.p < 0.05) before and after Losmapimod pre-treatment . **B,** heat map of gene expression of monocyte chemokines in normal and saline injected skin pre- and post-Losmapimod. **C,** Senescent fibroblasts were cultured with Losmapimod (3μM) for 3 or 24 hours, and supernatants were collected. CCL2 concentrations were assessed by cytometric bead array. Data assessed by paired t test * = p<0.05** = p<0.01 **** = p<0.000

|  | **Saline** | | | **VZV** | | **Air** | |
| --- | --- | --- | --- | --- | --- | --- | --- |
| **Donor characteristic** | **Young** | **Old** | **Old Improver** | **Young** | **Old** | **Young** | **Old** |
| **Age** | 25.0  (20.4-31.2) | 72  (66.9-79.3) | 70.0  (67.0-75.0) | 22.0  (19.0-28.0) | 71.5  (66.0-80.0) | 22.0  (20.0-24.0) | 75  (66.0-83.0) |
| **Gender** | 3 Male;  13 Female | 8 Male;  10 Female | 2 Male;  3 Female | 3 Male;  3 Female | 3 Male;  3 Female | 3 Male;  2 Female | 1 Male;  4 Female |
| **Clinical Score** | 4 | 1 | 5 | 5 | 1 | 4 | 1 |
| **Number of Donors** | 16 | 18 | 5 | 6 | 6 | 5 | 5 |

**Table 1: Skin donor characteristics divided according to age.**

Characteristics for all donors who donated skin for immunohistology and flow cytometric analysis for saline, VZV and air-injected skin. Data shown as Median with 10-90% Confidence intervals where appropriate.
